# Exchange between compartments regulates steady states and stochastic switching of a multisite phosphorylation network

**DOI:** 10.1101/2023.05.04.539393

**Authors:** Hannah N. Schmidt, Emily E. Leopin, Steven M. Abel

## Abstract

The phosphoregulation of proteins with multiple phosphorylation sites is governed by biochemical reaction networks that can exhibit multistable behavior. However, the behavior of such networks is typically studied in a single reaction volume, while cells are spatially organized into compartments that can exchange proteins. In this work, we use stochastic simulations to study the impact of compartmentalization on a two-site phosphorylation network. We characterize steady states and fluctuation-driven transitions between them as a function of the rate of protein exchange between two compartments. Surprisingly, the rate of stochastic switching between states depends nonmonotonically on the protein exchange rate, with the highest rate of switching occurring at intermediate exchange rates. At sufficiently small exchange rates, the state of the system and rate of switching are controlled largely by fluctuations in the balance of enzymes in each compartment. This leads to negatively correlated states in the compartments. For large exchange rates, the two compartments behave as a single effective compartment. However, when the compartmental volumes are unequal, the behavior differs from a single compartment with the same total volume. These results demonstrate that exchange of proteins between distinct compartments can regulate the emergent behavior of a common signaling motif.

## 1 Introduction

Compartmentalization is a key organizational principle in cell biology. The interiors of cells are organized in part by membrane-enclosed organelles and their more recently discovered membraneless counterparts [1, 2]. However, relatively little work has explored how signaling responses are regulated by compartmentalization and the exchange of proteins between compartments, especially in the context of membraneless organelles.

The dynamical and steady state behavior of intracellular signaling networks is governed by features including topology of the reaction network, kinetic parameters, and concentrations of proteins. Spatial organization and spatiotemporal correlations can also regulate the emergent behavior of signaling networks [3–7]. However, spatial effects are generally less well understood, as signaling networks are most commonly studied in well-mixed settings. It is likely that compartmentalization is a regulatory mechanism for some biochemical reaction networks because it connects mesoscale organization to dynamics of signaling networks [2, 8, 9].

Potential implications of compartmentalization are especially exciting in light of the vast and rapidly expanding field of biomolecular condensates. Membraneless organelles are condensed, liquid-like droplets containing proteins and other biomolecules. They are physically distinct from their surrounding environment, yet they are not separated by a phospholipid membrane and are able to exchange biomolecules with their surroundings [2, 10, 11]. While much emphasis has been placed on mechanisms of their formation, relatively little is known about how they regulate biomolecular processes in cells [9, 12, 13]. Recent work has shown that, even though protein levels in cells are noisy, liquid droplets can reduce the noise in protein concentration outside of the droplets [13, 14]. Additionally, Sang et al. recently engineered synthetic condensates that incorporate phosphorylation reactions [9]. The synthetic condensates exhibit increased kinase activity, with the potential to broaden kinase specificity when there are multiple potential substrates. Phase separation of liquid phases also appears to modulate membrane proximal immune-cell signaling [12], and it is exciting to consider the possibility of engineering spatial control in condensed systems. For example, macromolecular crowding was used to differentially organize components of cell-free protein synthesis, impacting the output of gene expression [15, 16].

The phosphoregulation of proteins is an important regulatory mechanism in cells. Kinases and phosphatases regulate the phosphorylation and dephosphorylation, respectively, of many proteins. Mitogen-activated protein kinase (MAPK) cascades are some of the most-studied signalling pathways in eukaryotic cells, propagating signals from the plasma membrane to the nucleus, enabling dynamic signal processing, and regulating many downstream processes [9, 17–20]. Other well-studied examples include networks involved in control of the cell cycle and in the phosphorylation of the cytoplasmic tail of the T-cell receptor complex [21].

In MAPK cascades, multisite phosphorylation provides a mechanism to control protein activity, leading to emergent behavior including ultrasensitivity and bistability [19, 22–25]. Multisite phosphorylation networks as simple as a single leaflet of the MAPK cascade can exhibit bistability when kinases and phosphatases bind in a distributive manner (i.e., they unbind after each enzymatic modification) [26]. Bistability describes the ability of a network to exist in one of two stable steady states, and it is a common feature of signaling networks that allows cells to make binary decisions [27]. More generally, networks can exhibit multistability, in which there are two or more stable steady states. Interestingly, Harrington et al. showed that allowing exchange of select species between two compartments can lead to the emergence of bistability in an otherwise monostable reaction network. They focused on the regulation of a substrate protein with a single phosphorylation site, demonstrating that compartmentalization expands the emergent behavior possible by a reaction network [20].

In multistable systems, fluctuations associated with stochastic reaction kinetics can lead to stochastic switching between steady states [28–30]. Behavior in such systems is typically characterized by long residence times in steady states, interspersed by rapid transitions between them [31, 32]. Spatial effects can shape the likelihood of stochastic switching in bistable networks [33, 34], but little is known about the impact of compartmentalization when components can exchange between compartments.

This paper explores the impact of compartmentalization on a network in which the phosphorylation states of substrate proteins are regulated by kinases and phosphatases. The network, detailed in the Methods, is motivated by a single leaflet of the MAPK network and by other networks involving the phosphoregulation of substrate proteins with multiple phosphorylation sites [3, 4]. We use stochastic, particle-based simulations to characterize steady states and stochastic switching between them. We first characterize behavior in a single compartment before systematically varying the exchange rate of particles between two compartments. For compartments of equal volume, we show that the exchange rate regulates steady states, correlation between the compartments, and the rate of stochastic switching between states. We then study the effects of particle exchange between compartments of different volumes. For both cases, the behavior is strongly influenced by fluctuations in the balance of enzymes at low and intermediate exchange rates. However, when exchange rates are large, compartments with unequal volumes give rise to different behavior than a single compartment with the same total volume. Taken together, our work emphasizes that compartmentalization and protein exchange can regulate the emergent behavior of common signaling motifs.

## 2 Methods

We use the Gillespie algorithm [35] to simulate the time evolution of a distributive, two-site phosphorylation network in two coupled compartments. The system is assumed to be well-mixed in each compartment, and proteins exchange between compartments with rate *k*_*E X*_. The network contains substrate proteins that can be phosphorylated at two sites: A kinase (*E*) catalyzes the phosphorylation of an unphosphorylated residue, and a phosphatase (*P*) catalyzes the dephosphorylation of a phosphorylated residue. Each substrate protein can have no sites (*S*_0_), one site (*S*_1_), or both sites (*S*_2_) phosphorylated. Distributive reaction networks require enzymes to unbind from the substrate they are modifying between each catalytic step [26, 36]. The following reactions specify the network in compartment *i* (*= A* or *B*):

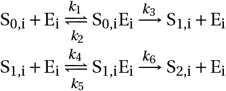

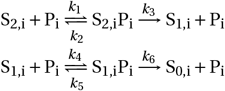

The compartments are coupled by the exchange of particles: Each protein or protein complex in compartment *A* transitions to compartment *B* with rate *k*_*E X*_, and vice versa. We use the kinetic parameters in Table 1, which are based on previous studies [3, 4]. The exchange rate is systematically varied to characterize the impact on steady states and the transitions between them.

**Table 1:** Kinetic parameters for the reaction network. Reactions associated with rates *k*_1_ – *k*_6_ take place in compartments *A* and *B* (subscripts are left off for clarity).

| Kinetic Parameter | Value | Related Reactions |
| --- | --- | --- |
| $k_1$ | $0.045 \mu\text{m}^3\text{s}^{-1}$ | $S_0 + E \rightarrow S_0E$<br>$S_2 + P \rightarrow S_2P$ |
| $k_2$ | $1.35 \text{ s}^{-1}$ | $S_0E \rightarrow S_0 + E$<br>$S_2P \rightarrow S_2 + P$ |
| $k_3$ | $1.5 \text{ s}^{-1}$ | $S_0E \rightarrow S_1 + E$<br>$S_2P \rightarrow S_1 + P$ |
| $k_4$ | $0.093 \mu\text{m}^3\text{s}^{-1}$ | $S_1 + E \rightarrow S_1E$<br>$S_1 + P \rightarrow S_1P$ |
| $k_5$ | $1.73 \text{ s}^{-1}$ | $S_1E \rightarrow S_1 + E$<br>$S_1P \rightarrow S_1 + P$ |
| $k_6$ | $15 \text{ s}^{-1}$ | $S_1E \rightarrow S_2 + E$<br>$S_1P \rightarrow S_0 + P$ |
| $k_{EX}$ | varied ($\text{s}^{-1}$) | $E_A \leftrightarrow E_B$<br>$P_A \leftrightarrow P_B$<br>$S_{x,A} \leftrightarrow S_{x,B}$<br>$S_{x,A}E_A \leftrightarrow S_{x,B}E_B$<br>$S_{x,A}P_A \leftrightarrow S_{x,B}P_B$ |

The system is initialized with 50 *S*_0_, 50 *S*_2_, 25 *E*, and 25 *P* in each compartment. The volume of the compartments is varied to characterize the effect of particle concentration. Because of the symmetry between kinase and phosphatase reactions, these initial conditions provide an unbiased initial state. For each condition studied, we generate 1,000 independent trajectories, each of which is 10,000 seconds in duration. The state of the system is recorded every 0.1 seconds, and the first 100 seconds of each trajectory are excluded from calculations to allow the system to reach steady state.

### 2.1 Detecting stochastic switches

When in a multistable regime, the system can stochastically switch between steady states due to intrinsic fluctuations in the system. To identify stochastic switches, we use a heuristic algorithm that analyzes time traces of the number of S_2_ molecules 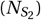 in each compartment. We identify switches by assessing when the moving average of 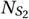 crosses a threshold value approximately equal to the value of unstable steady state identified from analysis of the deterministic equations near the critical point (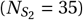. Specifically, we calculate a moving average with a time window of 10 s and classify a switching event in a compartment when the moving average crosses the threshold for at least 1 s. This reduces the overclassification of short-lived fluctuations as switching events. We use this information to determine the distribution of residence times in each steady state and to quantify the rate of stochastic switching.

### 2.2 Code availability

Simulation and analysis code used in this work is available here.

## 3 Results and discussion

### 3.1 An isolated compartment exhibits bistability at sufficiently high concentrations

We begin by analyzing the reaction network in a single, well-mixed compartment. We fix the number of proteins (100 substrate proteins, 25 kinases, and 25 phosphatases) and vary the volume. Figure 1 shows the number of fully phosphorylated substrate particles 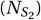 at steady state, as determined by stochastic simulations (red points) and numerical solutions of the ordinary differential equations associated with deterministic, mass-action kinetics (black lines). The deterministic solutions highlight a pitchfork bifurcation, with the system bistable at sufficiently small volumes and monostable at larger volumes. Below a critical volume (*≈* 0.46 *μ*m^3^), there are two stable steady states (solid lines) and one unstable steady state (dashed line) between them. Above the critical volume, there is a single stable steady state. For the stochastic simulations, we characterize the distribution of the number of *S*_2_ molecules. At sufficiently small volumes, the distribution is bimodal, and the location of each mode (red dot) is close to a stable deterministic solution. The distribution consists of a single mode in the monostable regime.

**Figure 1:**
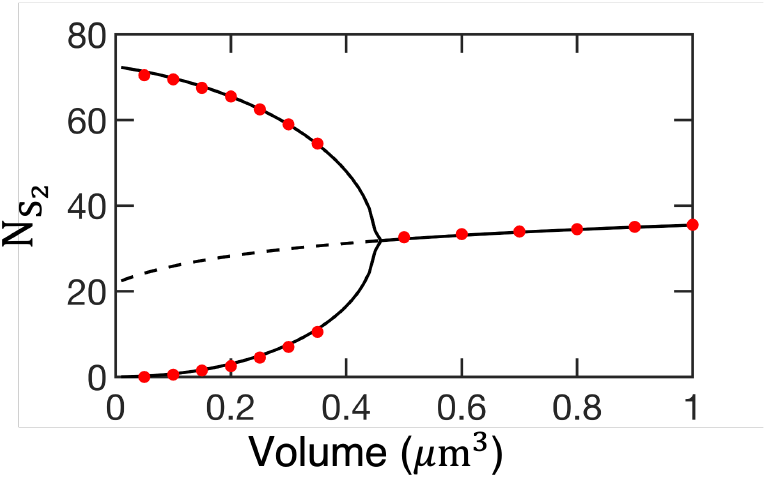
Number of *S*_2_ particles at steady state in a single, well-mixed volume, as determined by deterministic (black lines) and stochastic (red circles) methods. Solid lines denote stable steady states and the dashed line denotes an unstable steady state. Results from stochastic simulations correspond to the mode(s) of the distribution of 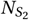.

In the bistable region, we refer to the state with more *S*_2_ particles as the *active state* and the state with fewer *S*_2_ particles as the *inactive state*. The behavior in Figure 1 is a consequence of changing concentration: The number of particles in the system is constant, so a smaller volume results in a larger concentration. For the two-site distributive reaction network, bistability arises from a sequestration effect when the number of substrate proteins exceeds the number of enzymes [4, 26]. When the system is in the active state 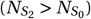, more phosphatases are typically bound than kinases. When a protein is dephosphorylated, the phosphatase unbinds, and there are more kinases than phosphatases available to bind to the now singly-phosphorylated protein. Thus, it is more likely to return to the fully phosphorylated state. An analogous argument holds for the inactive state 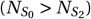, where the kinases are sequestered and there is an excess of phosphatases available to bind. Larger concentrations promote protein binding, which enhances the sequestration effect needed for bistability.

### 3.2 The exchange rate controls steady states and correlation between compartments

For the remainder of the paper, we consider a system with two compartments (*A* and *B*). Initially, we consider compartments of equal volume, with *V*_*A*_ *= V*_*B*_ *=* 0.16, 0.24, 0.32, and 0.4 *μ*m^3^. These volumes are within the bistable region identified in Figure 1 for a single compartment. Focusing on equal volumes allows us to examine the impact of particle exchange without confounding effects of different volumes. We later explore the effect of pairing compartments with *V*_*A*_*?= V*_*B*_ when, in the absence of exchange, compartment *A* would be bistable and compartment *B* would be monostable. When *k*_*E X*_ *=* 0, there is no particle exchange, and the compartments evolve independently. In this limit, each compartment is equivalent to a case considered in Section 3.1.

Figure 2 shows the time-dependence of 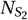 in compartments *A* and *B* for a portion of a single simulation trajectory. Switching events are identified independently in each compartment and are denoted by vertical lines. For this volume (*V*_*A*_ *= V*_*B*_ *=* 0.32 *μ*m^3^) and exchange rate (*k*_*E X*_ *=* 0.01 s^*−*1^), the switching events in the two compartments appear to occur at similar times. Further, the states of *A* and *B* appear to be anti-correlated: When compartment *A* is in an active state, compartment *B* tends to be in an inactive state, and vice versa.

**Figure 2:**
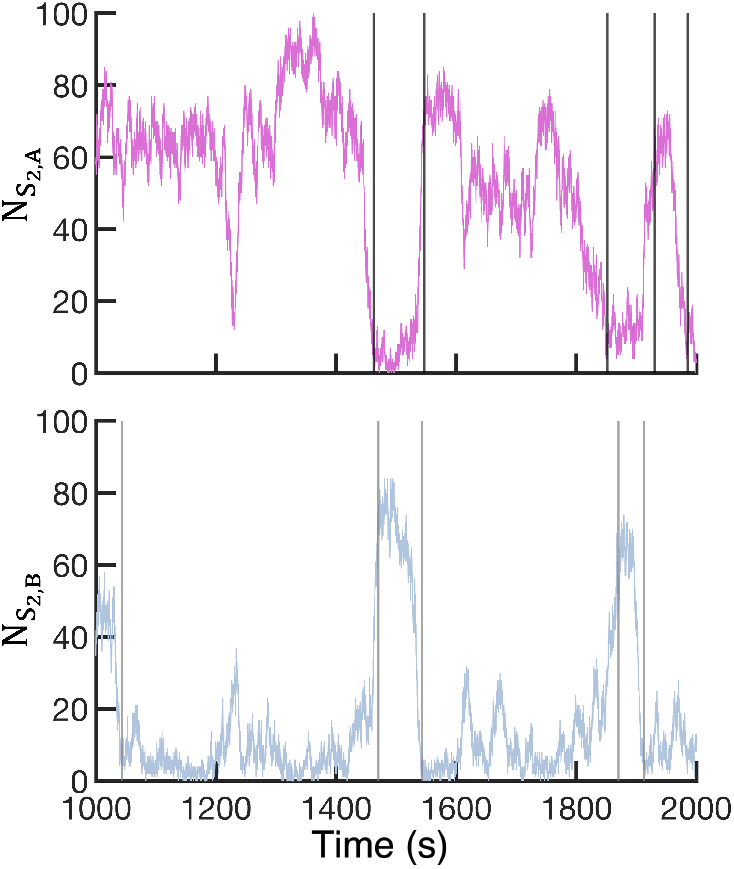
Time-dependent behavior of the number of *S*_2_ particles in compartments *A* (top) and *B* (bottom) for *V*_*A*_ *= V*_*B*_ *=* 0.32 *μ*m^3^ and *k*_*E X*_ *=* 0.01 s^*−*1^. A portion of a single trajectory is shown. Switching events are determined independently in each compartment and are denoted by vertical lines.

We show sample trajectories for additional exchange rates in Figure 3 (upper panel). For each exchange rate, each compartment exhibits two distinct states with stochastic switching between them. At *k*_*E X*_ *=* 0 s^*−*1^, the compartments behave independently of each other. When *k*_*E X*_ *=* 0.01 s^*−*1^ (see also Figure 2), the states in compartments *A* and *B* appear negatively correlated, with one compartment tending to be in the active state when the other is inactive. When *k*_*E X*_ *=* 1 s^*−*1^, the states of the two compartments are highly correlated, with both compartments having similar time dependence.

**Figure 3:**
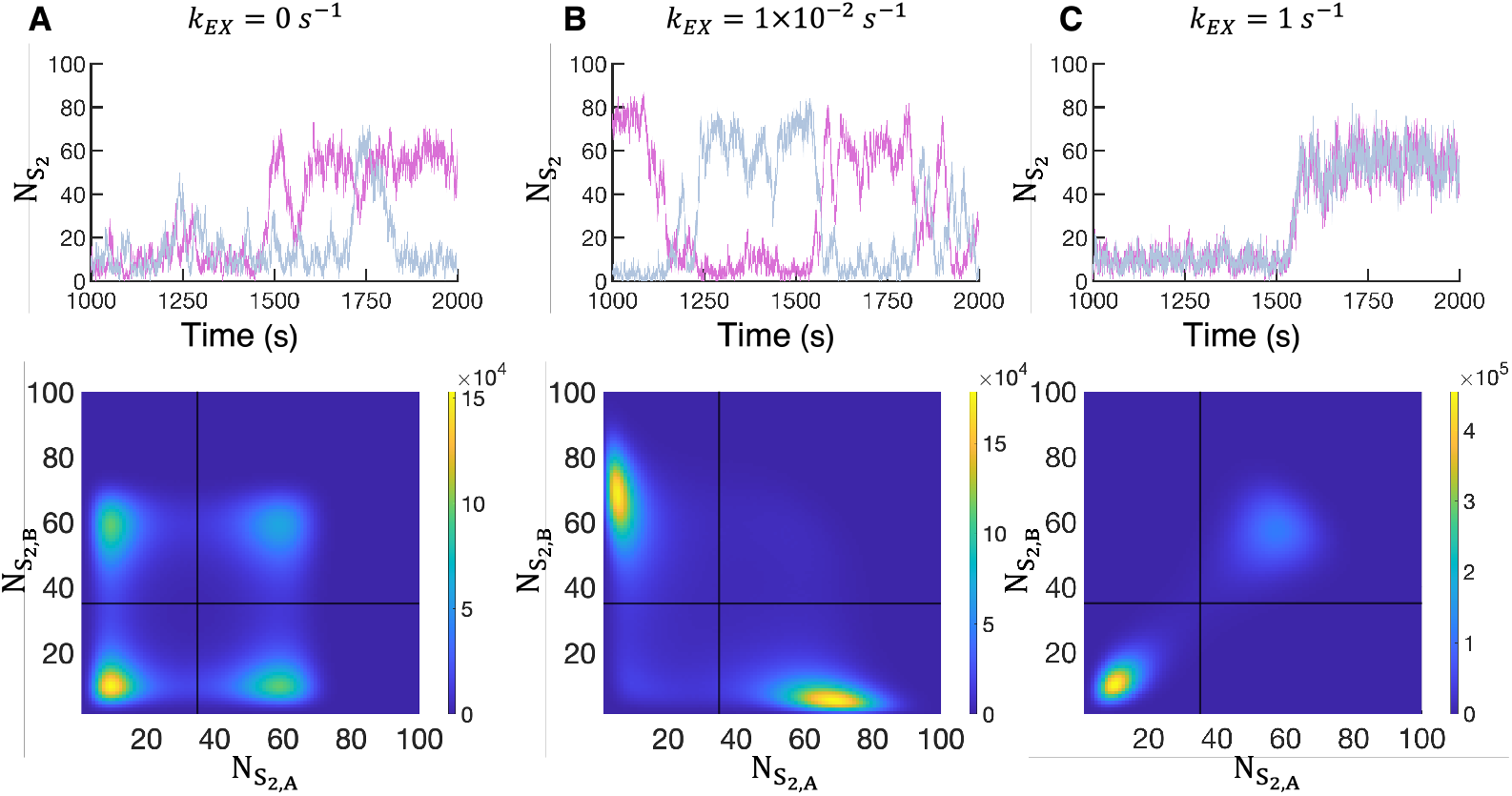
Behavior with *V*_*A*_ *= V*_*B*_ *=* 0.32 *μ*m^3^ at various exchange rates: (A) *k*_*E X*_ *=* 0 s^*−*1^, (B) *k*_*E X*_ *=* 0.01 s^*−*1^, and (C) *k*_*E X*_ *=* 1 s^*−*1^. The top panel shows the number of *S*_2_ particles in compartment *A* (fuchsia) and compartment *B* (blue) from part of a single trajectory. The bottom panel shows the distribution of *S*_2_ particles in compartments *A* and *B* sampled from 1, 000 independent trajectories. Horizontal and vertical black lines denote the threshold 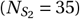 used to identify switches.

The lower panel in Figure 3 shows the simultaneous distribution of the number of *S*_2_ particles in compartments *A* and *B*. With *k*_*E X*_ *=* 0 s^*−*1^, the compartments evolve independently, and each undergoes independent stochastic switches between steady states. This is reflected in the four regions of high frequency in the two-dimensional distribution: The four states are associated with each compartment being either active or inactive, independent of the other. In contrast, the distribution for *k*_*E X*_ *=* 0.01 s^*−*1^ demonstrates the negative correlation suggested by the sample trajectories. The vast majority of the weight is associated with compartment *A* being active while compartment *B* is inactive, and vice versa. The active state is shifted to larger numbers of *S*_2_ molecules compared with *k*_*E X*_ *=* 0 s^*−*1^, and the shape of the distribution around the steady state also changes, with a larger range of 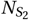 sampled. At the highest exchange rate, *k*_*E X*_ *=* 1 s^*−*1^, the distribution reflects the highly correlated time traces, with most of the weight associated with states in which both compartments are either active or inactive. Distributions obtained with other exchange rates are shown in Figure S1 and illustrate the transition between the negatively and positively correlated distributions.

To further characterize correlations between the states of the compartments, Figure 4A shows the fraction of time compartments *A* and *B* are in the same state. For all volumes, the fraction is *≈* 1/2 when *k*_*E X*_ *=* 0 s^*−*1^ (dashed line), indicating that the compartments are uncorrelated. Each volume exhibits a similar shape: At low exchange rates, the fraction of time in the same state is *<* 1/2, indicating that the states of the two compartments are negatively correlated. As the exchange rate increases, the system switches from negatively to positively correlated. At high exchange rates, the compartments are almost always in the same state. As the volume increases, the transition from negative to positive correlation between the compartments occurs at higher exchange rates.

**Figure 4:**
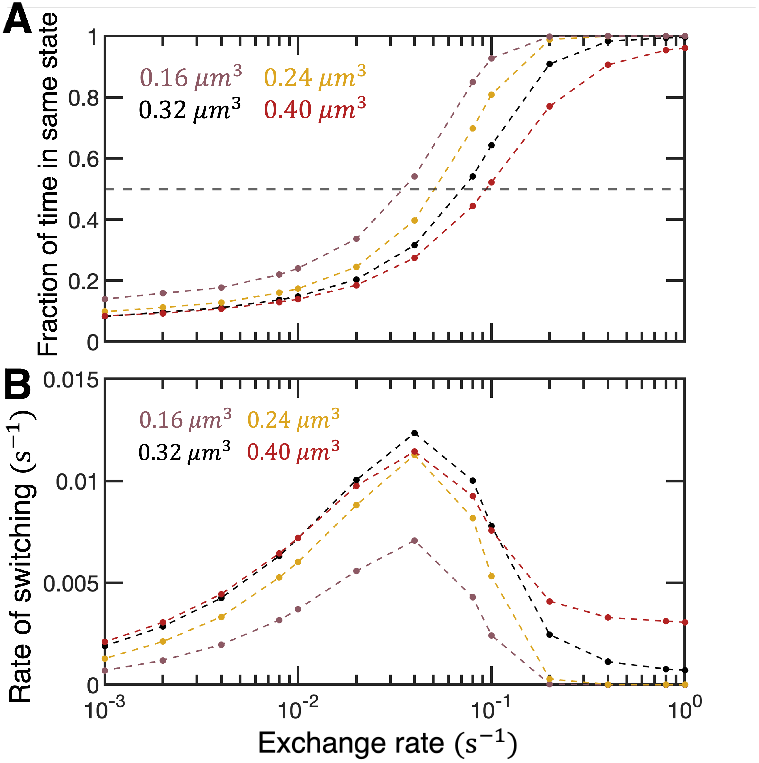
(A) Correlation between compartments characterized by the fraction of time the compartments are in the same state (active or inactive). The dashed line (*=* 1/2) is obtained when no exchange occurs and the compartments are uncorrelated. As the exchange rate increases, the compartments transition from negatively to positively correlated. (B) The rate of stochastic switching as a function of exchange rate. Each data point is obtained from 1,000 trajectories.

### 3.3 Stochastic switching is promoted at intermediate exchange rates

The sample trajectories shown in Figure 3 suggest a change in the frequency of stochastic switching as the exchange rate is varied. In Figure 4B, we show the rate of switching as a function of the exchange rate for different volumes. Note that the rate of switching is defined in terms of switching for a single compartment. This facilitates comparison to the single-compartment case and to cases with compartments of different volumes.

Figure 4B reveals that the rate of stochastic switching is a nonmonotonic function of the exchange rate, and that the maximum rate of switching occurs at intermediate exchange rates. At low exchange rates, the rate of switching increases with increasing exchange rate. The rate of switching then peaks at intermediate exchange rates, before rapidly falling to a plateau at higher exchange rates. Smaller volumes exhibit less frequent stochastic switching. However, the differences are modest at low and intermediate exchange rates when compared with isolated compartments. With *k*_*E X*_ *=* 10^*−*3^ s^*−*1^, the largest volume switches *≈* 3 times more frequently than the smallest volume. In contrast, when *k*_*E X*_ *=* 0 s^*−*1^, the largest volume switches *≈* 8600 times more frequently than the smallest volume.

For large exchange rates, the differences in the switching rates between different volumes are more pronounced. In this regime, the switching rates for the smallest and largest volumes vary by over two orders of magnitude. Physically, the two compartments behave like a well-mixed system with a larger effective volume (*V = V*_*A*_ *+V*_*B*_). To test this, we consider a single compartment with total volume *V* and the same total number of particles. Figure S2 shows that, when the exchange rate is large, the distribution of the total number of *S*_2_ particles in both compartments is almost indistinguishable from the distribution of the single compartment. Because of the larger number of particles at the same concentration, the impact of intrinsic fluctuations is reduced, thus suppressing stochastic switching when the exchange rate is large [28, 37].

### 3.4 Fluctuations in the balance of enzymes influence steady states at low and intermediate exchange rates

Figure 4B shows that the rate of particle exchange influences the frequency of switching. Further, Figures 3 and S1 reveal, at low and intermediate exchange rates, a negative correlation between the compartments, an increase in the value of 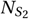 in the active state, and a broadening of the distribution associated with the active state. Taken together, these results suggest a mechanism for stochastic switching that is influenced by the balance of enzymes in the compartments and the timescale of their fluctuations.

To test this, we characterize Δ_*EP*_, the difference in the total number of kinases and phosphatases in a compartment (the total includes both free and bound enzymes). Figures 5A and B show sample trajectories in which the time dependence of Δ_*EP*_ is plotted with 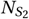 in the same compartment. With *k*_*E X*_ *=* 0.01 s^*−*1^, there is a strong correlation between fluctuations of Δ_*EP*_ and 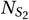 : When Δ_*EP*_ *>* 0, the compartment is likely to be in an active state, and when Δ_*EP*_ *<* 0, the compartment is likely to be in an inactive state. In contrast, with *k*_*E X*_ *=* 1 s^*−*1^, the fluctuations of Δ_*EP*_ occur on shorter timescales and appear uncorrelated with the state of the compartment.

**Figure 5:**
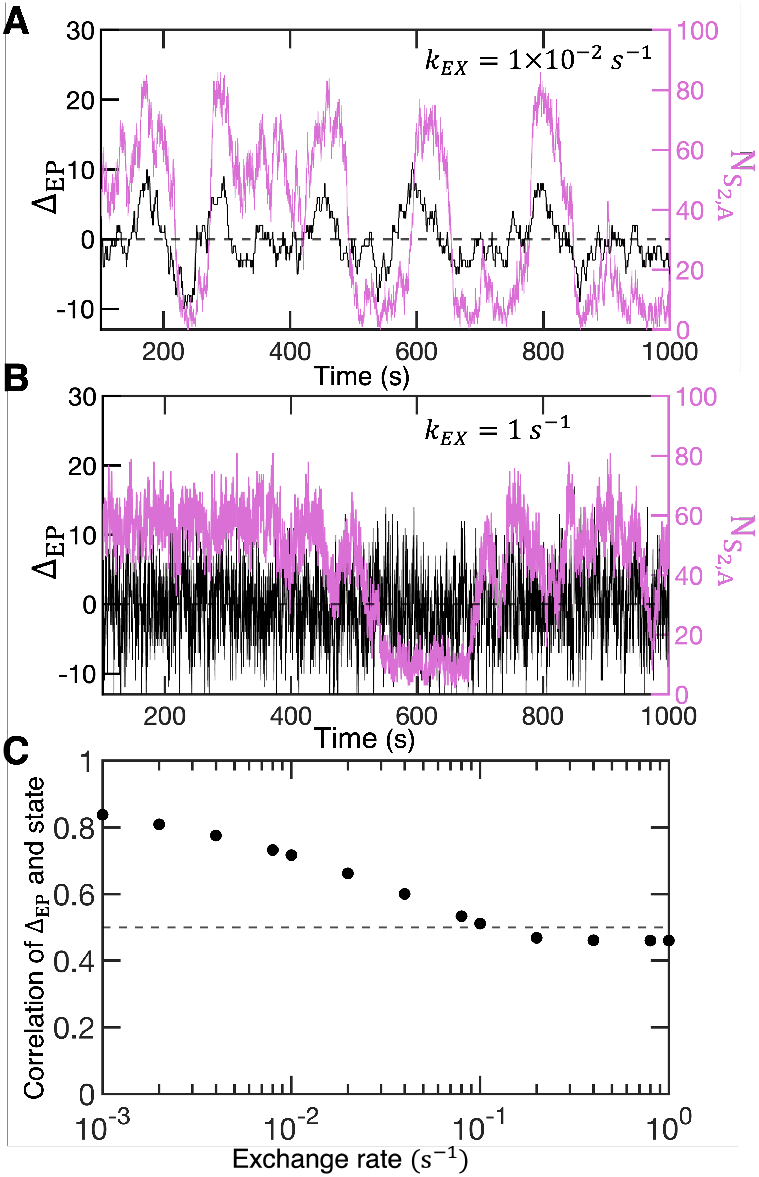
(A) Part of a trajectory showing 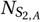,*A* and the difference in the number of kinases and phosphatases (Δ_*EP*_) in compartment *A* as a function of time for *k*_*E X*_ *=* 1 *×* 10^*−*2^ s^*−*1^ and *V*_*A*_ *= V*_*B*_ *=* 0.32 *μ*m^3^. (B) Analogous results with *k*_*E X*_ *=* 1 s^*−*1^. (C) The fraction of time a compartment is in the active state with an excess of kinase or the compartment is in an inactive state with an excess of phosphatase (1,000 trajectories for each exchange rate). As the exchange rate between compartments increases, the state of the system and Δ_*EP*_ become uncorrelated.

To further explore the relationship between Δ_*EP*_ and the state of the compartment, we determine the fraction of time a compartment is (i) in the active state with an excess of kinase (Δ_*EP*_ *>* 0) or (ii) in an inactive state with an excess of phosphatase (Δ_*EP*_ *<* 0). Figure 5C shows that the sign of Δ_*EP*_ is strongly correlated with the state of the compartment at low exchange rates. The correlation decreases with increasing exchange rate and becomes uncorrelated at higher exchange rates (the value is slightly less than 1/2 because we do not consider Δ_*EP*_ *=* 0). Thus, at low and intermediate exchange rates, when there are more phosphatases than kinases in a compartment, the compartment tends to be in the inactive state; similarly, when there are more kinases than phosphatases, the compartment tends to be in the active state. Changes in the balance of enzymes influence the transitions between active and inactive states. Other dynamical variables considered do not show strong correlation with the state of the system. For example, it is plausible that fluctuations in the total number of substrates in a compartment could bias the state of the network due to sequestration effects. We examine this variable in Figure S3, which shows no correlation with the state of the network across all exchange rates.

In Figure 6, we further characterize the relation between the balance of enzymes and the state of the compartment. Here we show the conditional distribution of 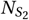 in a compartment given a specific value of Δ_*EP*_. For *k*_*E X*_ *=* 0 s^*−*1^, there is no enzyme exchange and Δ_*EP*_ *=* 0 for the entire simulation. This case gives the steady state distribution of 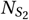 in the absence of exchange. With *k*_*E X*_ *=* 0.01 s^*−*1^, the distribution with Δ_*EP*_ *=* 0 is similar to the case with no exchange, with the distribution slightly broadened about each mode. As Δ_*EP*_ increases, indicating more kinases than phosphatases, the weight associated with the active state increases and the mode shifts to larger values. With Δ_*EP*_ *=* 8, there is only a single mode, indicating that the imbalance in enzymes biases the system to a single active state. Similarly, with Δ_*EP*_ *<* 0, the excess of phosphatases biases the system to an inactive state. In this regime, as Δ_*EP*_ decreases, the weight associated with the inactive state increases and the mode shifts to smaller values. In contrast, with fast exchange between compartments (*k*_*E X*_ *=* 1 s^*−*1^), the distributions show almost no dependence on Δ_*EP*_ in the range considered. This is consistent with the results of Figure 5 showing no correlation between enzyme fluctuations and the state of the system.

**Figure 6:**
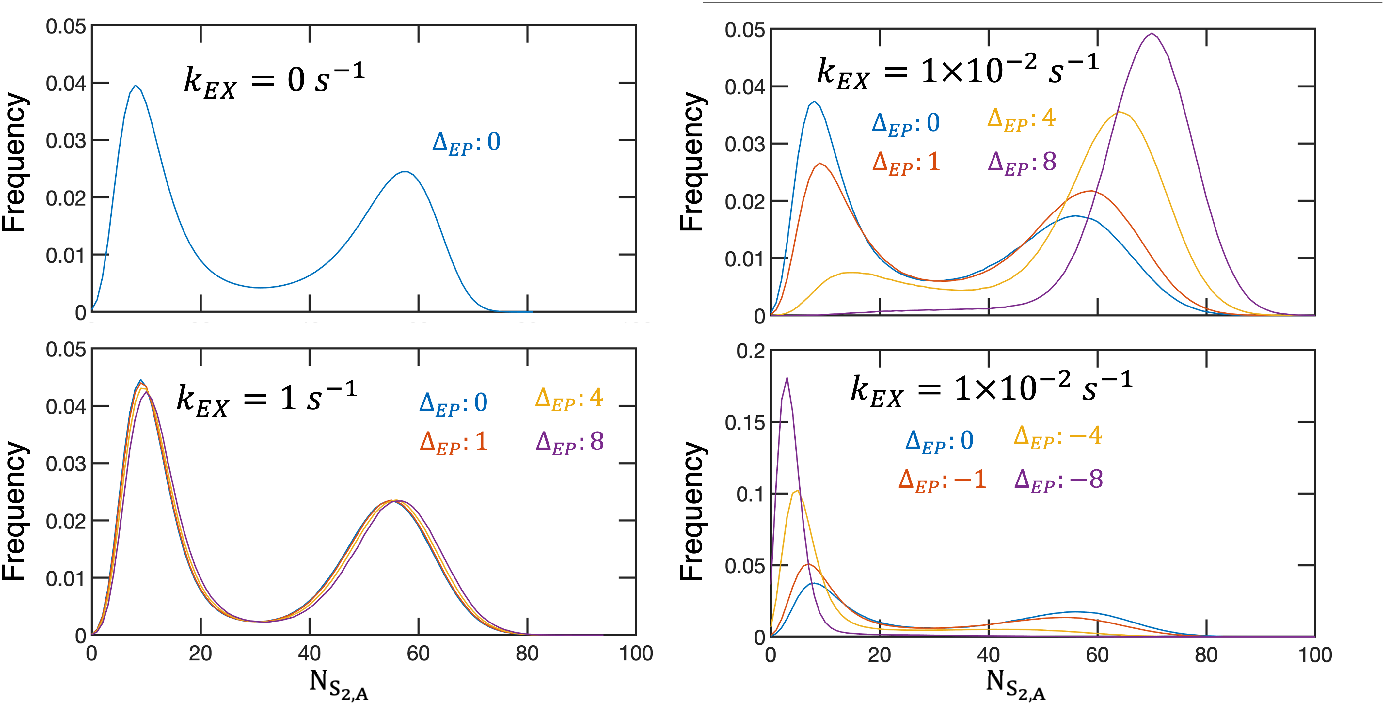
Conditional distribution of 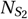 given specific values of Δ_*EP*_. Three exchange rates are shown with *V*_*A*_ *= V*_*B*_ *=* 0.32 *μ*m^3^. Results are obtained from 1,000 independent trajectories.

The bias introduced by imbalances in the enzymes leads to the negative correlation between the state of each compartment at low and intermediate exchange rates (Figure 4A). Because of particle conservation, an excess of kinases in compartment *A* implies an excess of phosphatases in compartment *B*, thus leading to anticorrelated states.

### 3.5 Coupling compartments of different sizes

Our results thus far have emphasized the importance of particle exchange between compartments when the average concentration is equal in each. In biological systems, it is common to encounter compartments with different effective concentrations of proteins. In this section, we consider two compartments with the same numbers of particles as above but with different volumes: *V*_*A*_ *=* 0.32 *μ*m^3^ and *V*_*B*_ *=* 0.8 or 10 *μ*m^3^. With no exchange, compartment *A* is in the bistable regime and compartment *B* is in the monostable regime (Fig. 1).

Figure 7 shows, for *V*_*A*_ *=* 0.32 *μ*m^3^ and *V*_*B*_ *=* 0.8 *μ*m^3^, sample trajectories (upper panel) and the distribution of the number of *S*_2_ particles in compartments *A* and *B* (lower panel) for three exchange rates. With *k*_*E X*_ = 0 s^*−*1^, the results reflect the bistable behavior in compartment *A* and monostable behavior in compartment *B*. With *k*_*E X*_ *=* 0.01 s^*−*1^, there remain two steady states, as can be seen in the distribution. However, compared to the case of no exchange, they are shifted, broadened, and negatively correlated. In compartment *A*, the active state is shifted to larger values of 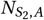 _,*A*_ and the inactive state is shifted to smaller values. Additionally, the distribution around each steady state exhibits negative correlation between 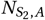 _,*A*_ and 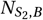 _,*B*_. This behavior is conceptually similar to that with *V*_*A*_ *= V*_*B*_, suggesting that the balance of kinases and phosphatases is again important in controlling the state of the system. With fast exchange (*k*_*E X*_ *=* 1 s^*−*1^), the system is monostable, as indicated by the single peak in the distribution. The states of compartments *A* and *B* are highly correlated, as seen in the trajectory and in the strong positive correlation in the distribution. The fluctuations are large, leading to a broad distribution of 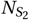 in each compartment. Distributions for additional exchange rates are shown in Figure S5A.

**Figure 7:**
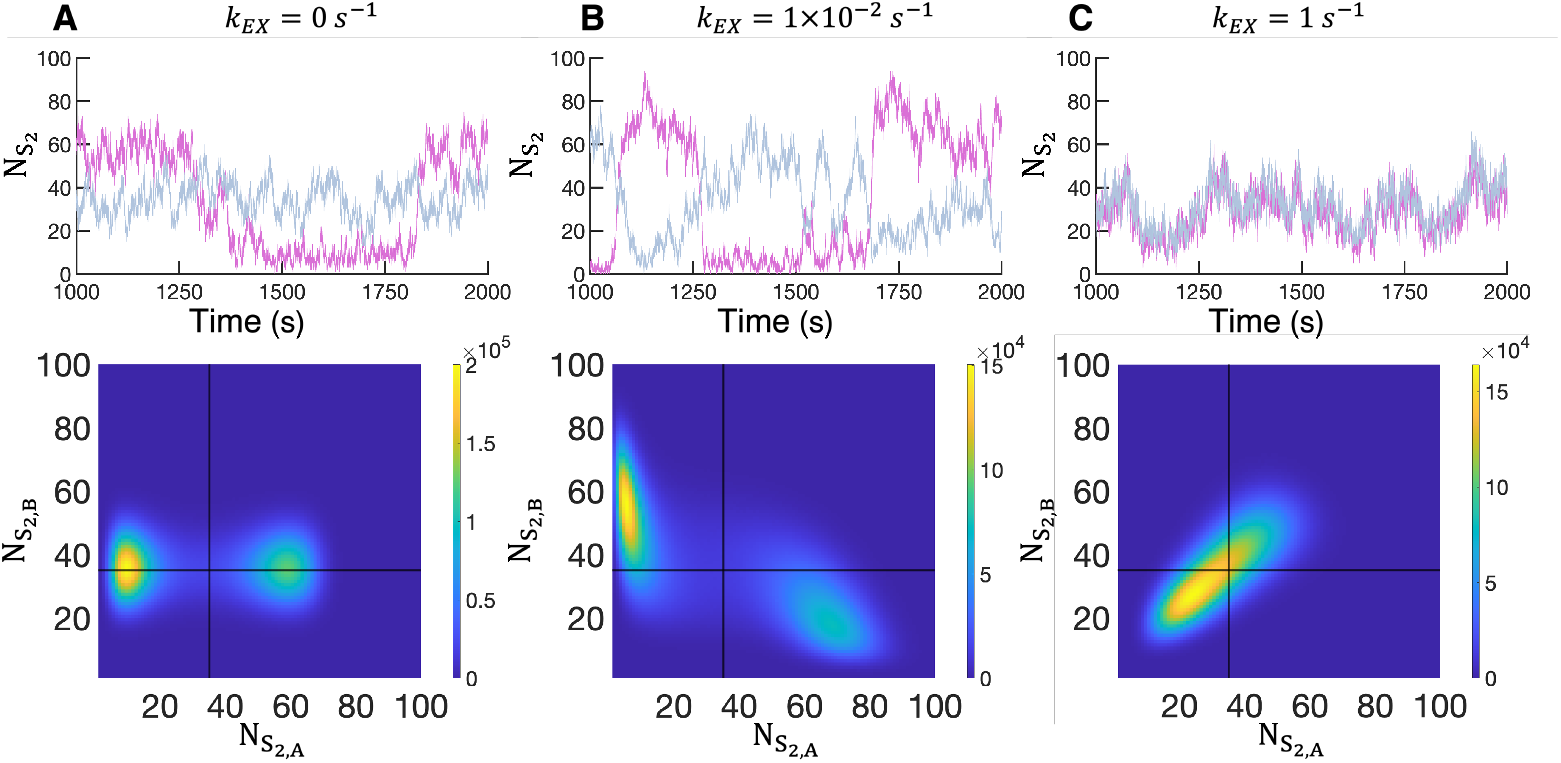
Behavior with *V*_*A*_ *=* 0.32 *μ*m^3^ and *V*_*B*_ *=* 0.8 *μ*m^3^ at various exchange rates: (A) *k*_*E X*_ *=* 0 s^*−*1^, (B) *k*_*E X*_ *=* 0.01 s^*−*1^, and (C) *k*_*E X*_ *=* 1 s^*−*1^. The top panel shows the number of *S*_2_ particles in compartment *A* (fuchsia) and compartment *B* (blue) from part of a single trajectory. The bottom panel shows the distribution of *S*_2_ particles in compartments *A* and *B* sampled from 1, 000 independent trajectories.

The behavior of the system with *V*_*A*_ *=* 0.32 *μ*m^3^ and *V*_*B*_ *=* 10 *μ*m^3^ is similar (Figures S5B), except that at low and intermediate exchange rates, the distribution of 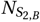,*B* is narrower. In this regime of exchange rates, enzyme imbalances remain important. However, because of the large volume of compartment *B*, enzymes are less likely to bind to substrates, thus reducing the effect of fluctuations in the balance of kinases and phosphatases in the larger compartment.

We further characterize the rate of switching in compartment *A* for exchange rates at which it exhibits bistable behavior. These results are shown in Figure 8 for *V*_*A*_ *=* 0.32 *μ*m^3^ and *V*_*B*_ *=* 0.32, 0.8, and 10 *μ*m^3^. In the slow-exchange regime, the behavior of the three cases is essentially indistinguishable. At intermediate exchange rates, coupling to larger compartments causes the peak of the switching rate to shift to modestly higher exchange rates, but the shape of the response is qualitatively similar. The rate of switching starts to decrease after the peak, but the system becomes monostable. These results indicate that the switching behavior of a compartment at low exchange rates is largely dependent on fluctuations in particle numbers due to exchange between the compartments, and not directly on the state of the other compartment.

**Figure 8:**
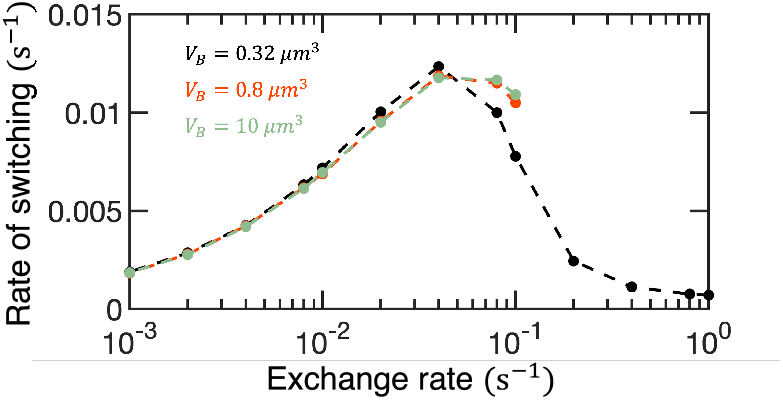
The rate of stochastic switching in compartment *A* as a function of exchange rate when *V*_*A*_ *=* 0.32 *μ*m^3^ and *V*_*B*_ *=* 0.32, 0.8, or 10 *μ*m^3^. Results with *V*_*B*_ *=* 0.32 *μ*m^3^ are also shown in Figure 4. With *V*_*B*_ *=* 0.8 and 10 *μ*m^3^, the system is monostable for *k*_*E X*_ ≳ 0.2 s^*−*1^, and hence switching times are not shown in this regime. Each switching rate is obtained from 1,000 independent trajectories.

For large exchange rates, the states of two compartments are highly correlated. Particles rapidly switch between compartments and thus effectively sample volume *V = V*_*A*_ *+ V*_*B*_ over short time scales. To this end, we compare the results with two compartments at large exchange rates (*k*_*E X*_ *=* 1, 10, and 100 s^*−*1^) with the equivalent single-compartment system of volume *V* (containing the same total number of each protein). Figures 9A and B compare the distribution of the total number *S*_2_ particles in two compartments with the number of *S*_2_ particles in the equivalent single compartment. The distributions are unimodal but markedly different in shape. With two compartments, the distribution is broader, indicating that the interplay between the two compartments is more complex than simply combining the two volumes. These results are in contrast with Figure S4, which shows that fast exchange with *V*_*A*_ *= V*_*B*_ leads to a steady state that is nearly indistinguishable from the equivalent single compartment.

**Figure 9:**
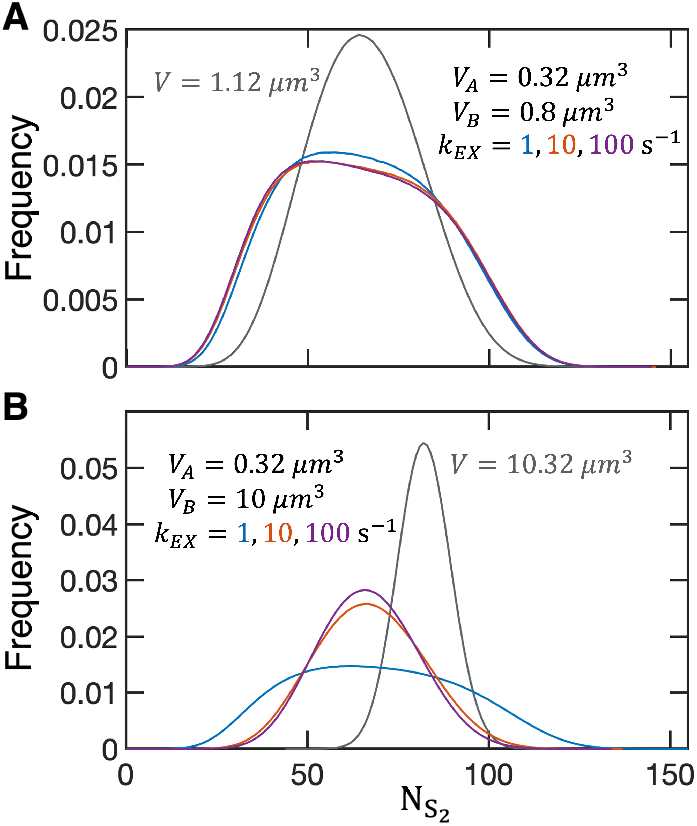
Distribution of 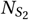 for two compartments and the equivalent single-compartment system with total volume *V = V*_*A*_ *+ V*_*B*_. The total number of particles is equal in each system. The two-compartment system is considered in the regime of rapid exchange, with *k*_*E X*_ *=* 1, 10, and 100 s^*−*1^. To facilitate comparison, the distribution for the two-compartment system corresponds to the total number of *S*_2_ particles in both compartments. Results are shown for (A) *V*_*A*_ *=* 0.32 *μ*m^3^, *V*_*B*_ *=* 0.8 *μ*m^3^ and (B) *V*_*A*_ *=* 0.32 *μ*m^3^, *V*_*B*_ *=* 10 *μ*m^3^.

## 4 Conclusions

Compartmentalization is a cornerstone of cell biology. Membrane-enclosed organelles such as the nucleus exchange proteins with the cytoplasm, with translocation of MAP kinases being one prominent example [7, 38]. Proteins can be recruited from the cytoplasm to the plasma membrane, creating two effective compartments [8, 39]. It is especially exciting to consider compartmentalization in the context of membraneless organelles, in which liquid-liquid phase separation leads to distinct domains that can be found in the cytoplasm, the nucleus, and the plasma membrane [1, 11, 40]. In all cases, proteins can dynamically exchange between different compartments and thus potentially impact signal transduction and other cellular processes.

Relatively little is known about the effects of compartmentalization and the exchange of proteins on the emergent behavior of signaling networks. To gain insight, we studied a common signaling motif describing the phosphoregulation of substrate proteins by kinases and phosphatases. The network and parameters were motivated by a leaflet of the MAPK pathway, which can exhibit bistability due to the sequestration of enzymes at sufficiently high concentrations.

Our key results are highlighted in Figure 4, which reveals a nonmonotonic dependence of the stochastic switching rate on the exchange rate of proteins. Surprisingly, the largest rate of stochastic switching occurs at intermediate exchange rates, revealing a nontrivial impact of protein exchange on emergent behavior of the reaction network. More detailed analysis revealed the importance of fluctuations in the balance of kinases and phosphatases in the compartments, which impacted the nature of the steady states and the stochastic switching between them.

At low exchange rates, the states of the compartments were negatively correlated, and the rate of switching was largely decoupled from the switching rate of isolated compartments. In this regime, the rate of switching increased with increasing exchange rate. These results are a consequence of fluctuations in enzyme numbers: When one compartment has an excess of kinases (Δ_*EP*_ *>* 0), the other has an excess of phosphatases (Δ_*EP*_ *<* 0). An excess of kinases promotes the active state, while an excess of phosphatases promotes the inactive state. Further, increasing the exchange rate decreases the average time during which Δ_*EP*_ remains positive or negative. Increasing the exchange rate promoted stochastic switching until a maximum at intermediate exchange rates. Here, the switching rate far exceeded that of an isolated compartment. However, at larger exchange rates, fluctuations in the balance of enzymes occur faster than the response time of the network. Thus, in this regime, the two-compartment system behaves more like a single, well-mixed volume. Interestingly, when the two compartments had different effective concentrations of particles, the coupled behavior was markedly different than the behavior of a single, well-mixed volume with the same overall concentration. This suggests a way to modulate the behavior of signaling networks using compartmentalization even when the exchange between compartments occurs on fast timescales.

These results demonstrate that compartmentalization and protein exchange can act as regulatory mechanisms for signaling networks. They also highlight the importance of the balance of kinases and phosphatases for a phosphoregulation network. When small numbers of proteins are involved, stochastic fluctuations in the balance of enzymes can drive stochastic switching between steady states in a multistable reaction network. Interesting directions for future studies include characterizing transition pathways between steady states in compartmentalized systems and studying signal propagation and regulation with multiple compartments. Recent work demonstrating engineered synthetic condensates [9] also opens exciting new avenues to use compartmentalization to control networks of biomolecular interactions. Computational approaches will likely provide a useful tool for understanding and designing responses in such systems.

