## Supplemental figures for "Exchange between compartments regulates steady states and stochastic switching of a multisite phosphorylation network"

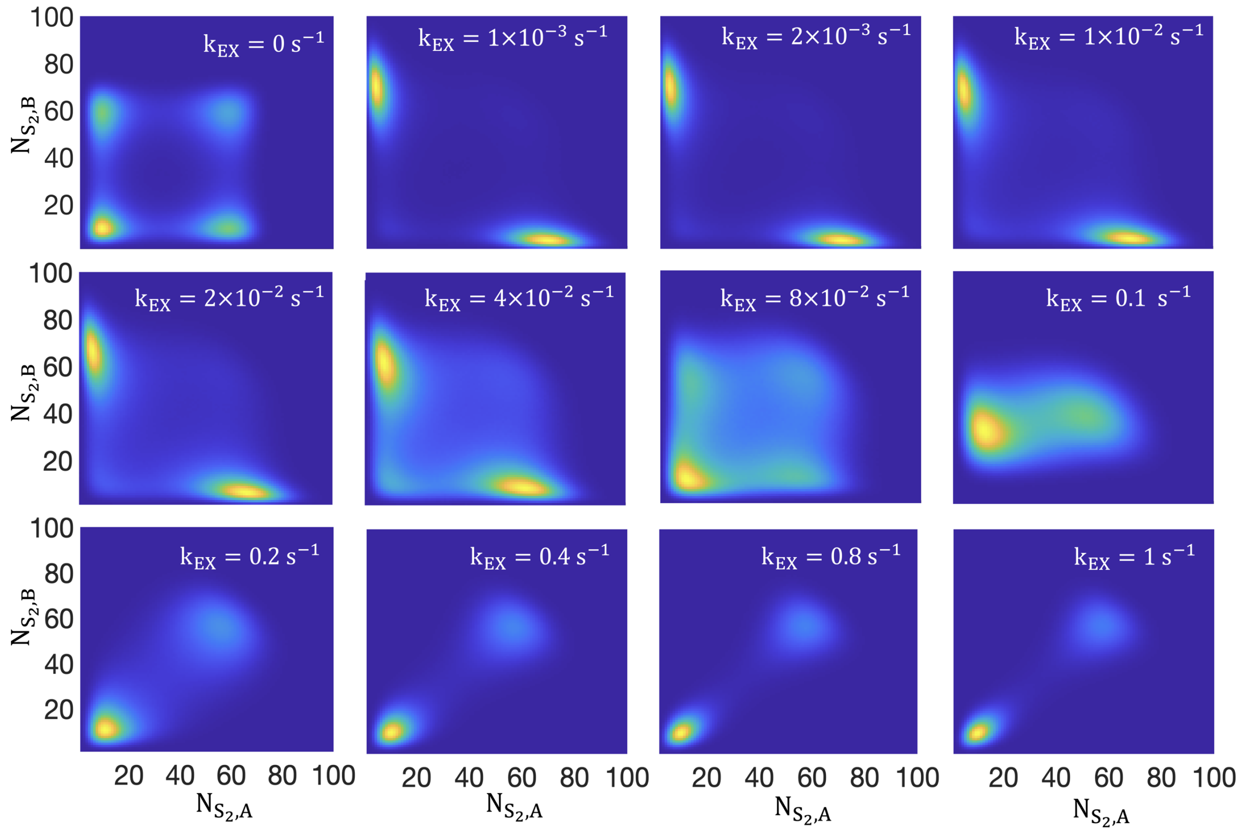


Figure S1: Distribution of $S_2$ particles in compartments A and B with $V_A = V_B = 0.32\ \mu m^3$. Select exchange rates are shown to highlight qualitative changes in the distribution. Each distribution is constructed from 1,000 independent trajectories.

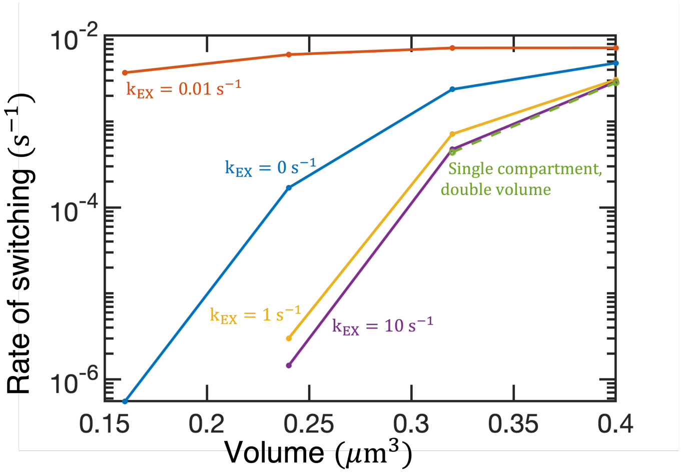


Figure S2: Rate of stochastic switching as a function of volume. Results are shown for two compartments with $k_{EX}$ = 0, 0.01, 1, and 10 $s^{-1}$ (solid lines, volume indicates that of a single compartment). The dashed line (green) shows the rate of stochastic switching for a single compartment with double the volume but the same concentration of particles. No switching events were observed for $V_A = V_B = 0.16$ $\mu m^3$ for $k_{EX} = 1$ and 10 $s^{-1}$ or for the single volume at the two smallest volumes considered.

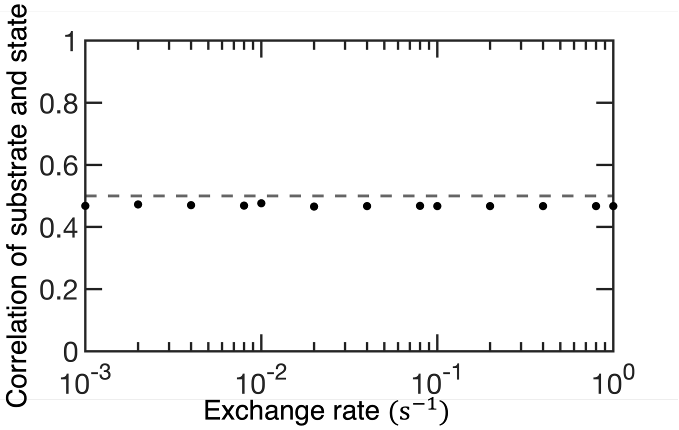


Figure S3: The fraction of time a compartment is in the active state with an excess in the total number of substrate particles (> 100) or in the inactive state with a deficiency in the total number of substrate particles (< 100). The two variables are uncorrelated across all exchange rates; the value lies below 0.5 (dashed line) because we do not consider the case with equal numbers of substrates. The lack of correlation is in contrast with $\Delta_{EP}$, which shows strong correlation with the state of the system at low values of the exchange rate.

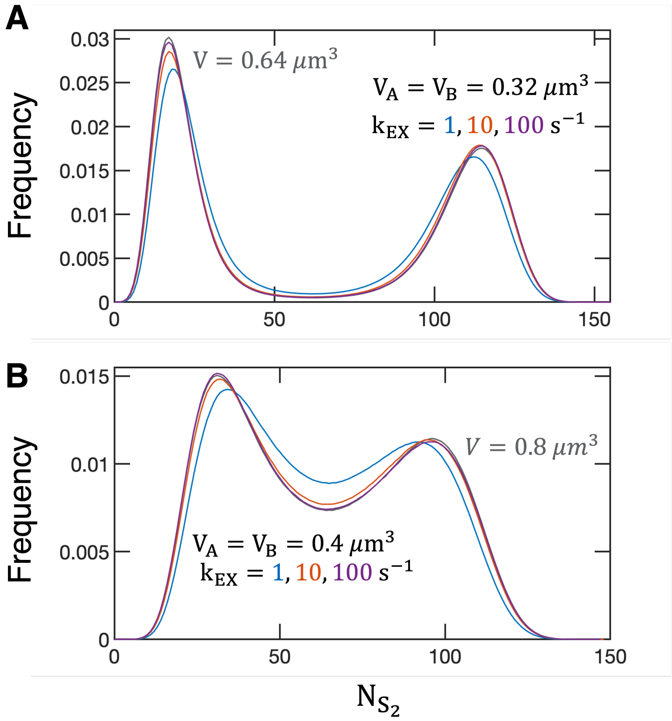


Figure S4: Distribution of $N_{S_2}$ for two compartments and the equivalent single-compartment system with total volume $V = V_A + V_B$. The total number of particles is equal in each system. The two-compartment system is considered in the regime of rapid exchange, with $k_{EX} = 1$, 10, and 100 $s^{-1}$. To facilitate comparison, the distribution for the two-compartment system corresponds to the total number of $S_2$ particles in both compartments. Results are shown for (A) $V_A = V_B = 0.32\ \mu m^3$ and (B) $V_A = V_B = 0.4\ \mu m^3$.

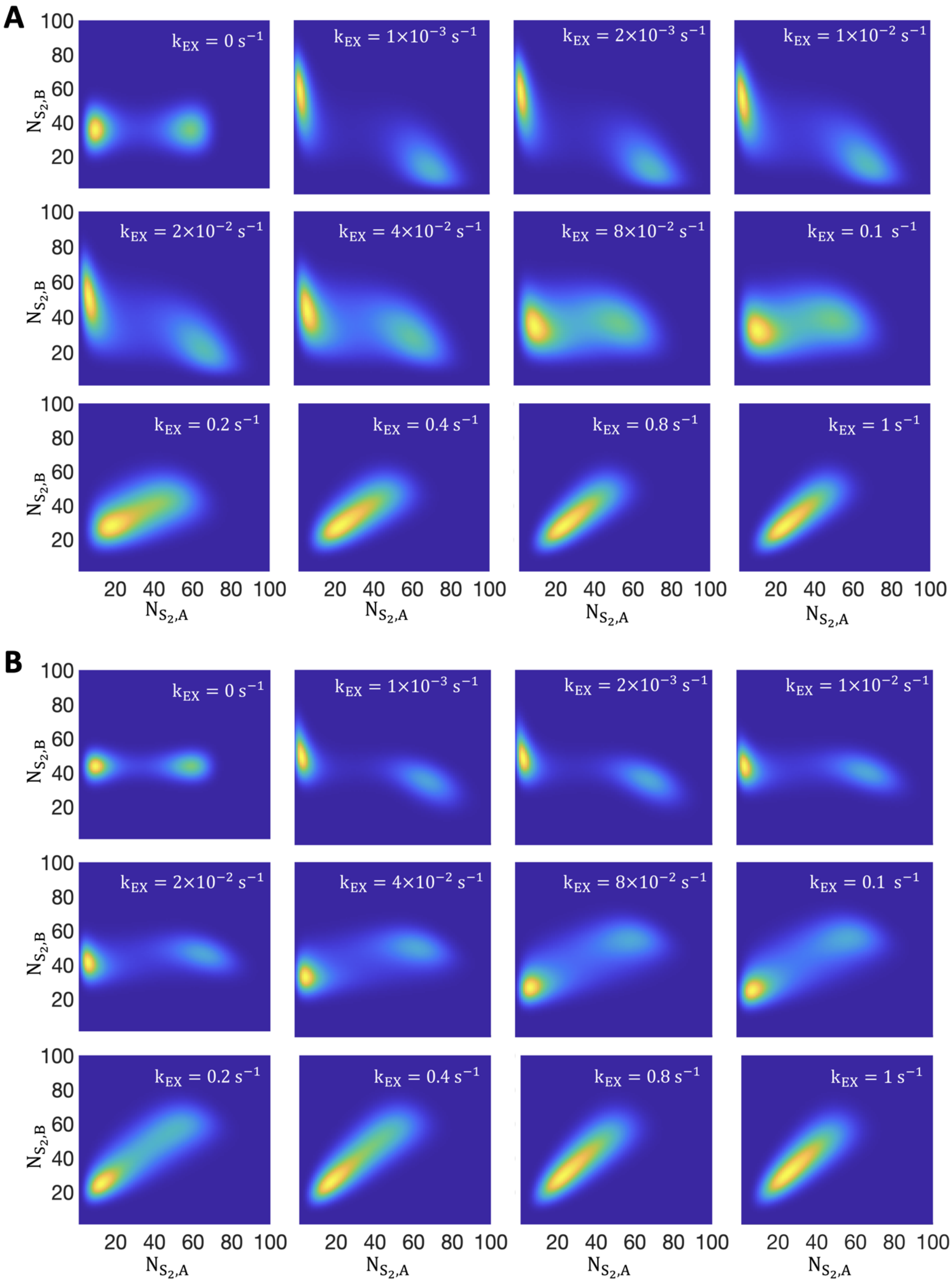


Figure S5: Distribution of $S_2$ particles in compartments A and B with with (A) $V_A = 0.32\ \mu\text{m}^3$ and $V_B = 0.8\ \mu\text{m}^3$ and (B) $V_A = 0.32\ \mu\text{m}^3$ and $V_B = 10\ \mu\text{m}^3$. Select exchange rates are shown to highlight qualitative changes in the distribution. Each distribution is constructed from 1,000 independent trajectories.
